# SARM1 deficiency, which prevents Wallerian degeneration, upregulates XAF1 and accelerates prion disease

**DOI:** 10.1101/485052

**Authors:** Caihong Zhu, Bei Li, Karl Frontzek, Yingjun Liu, Adriano Aguzzi

## Abstract

SARM1 (sterile α and HEAT/armadillo motifs containing protein) is a member of the MyD88 (myeloid differentiation primary response gene 88) family which mediates innate immune responses. Because inactivation of SARM1 prevents various forms of axonal degeneration, we tested whether it might protect against prion-induced neurotoxicity. Instead, we found that SARM1 deficiency exacerbates the progression of prion pathogenesis. This deleterious effect was not due to SARM1-dependent modulation of prion-induced neuroinflammation, since microglial activation, astrogliosis and brain cytokine profiles were not altered by SARM1 deficiency. Whole-transcriptome analyses indicated that SARM1 deficiency led to strong, selective overexpression of the pro-apoptotic gene XAF1 (X-linked inhibitor of apoptosis-associated factor 1). Consequently, the activity of proapoptotic caspases and neuronal death were enhanced in prion-infected *SARM1*^−/−^ mice. These results point to an unexpected function of SARM1 as a regulator of prion-induced neurodegeneration, and suggest that XAF1 might constitute a therapeutic target in prion disease.

## Introduction

SARM1 (sterile α and HEAT/armadillo motifs containing protein), a member of MyD88 (myeloid differentiation primary response gene 88) family (MyD88-5), is a highly conserved cytosolic adaptor molecule that negatively regulates TRIF (TIR domain containing adaptor inducing interferon-β) mediated Toll-like receptor (TLR) signaling in mammalian cell cultures (Carty et al., 2006). SARM1 differs from all other MyD88 family members in its structure, expression pattern and functions. SARM1 contains N-terminal HEAT/armadillo repeats and two sterile α motifs (SAM), followed by a C-terminal Toll-Interleukin 1 receptor (TIR) domain (Mink et al., 2001). SARM1 is expressed primarily by neurons in the mouse brain(Kim et al., 2007). In olfactory neurons of C. elegans, it regulates olfactory receptor expression (Chuang and Bargmann, 2005). By decreasing the association of JNK3 with mitochondria, SARM1 deficiency protects neurons from death due to oxygen and glucose deprivation (Kim et al., 2007). Similarly, *SARM1*^−/−^ mice also show protection from neuronal injury induced by vesicular stomatitis virus (VSV) and La Crosse virus (LACV) (Hou et al., 2013; Mukherjee et al., 2013). On the other hand, SARM1 restricts neurotropic West Nile Virus (WNV) infection by modulating microglia activation and cytokine production (Szretter et al., 2009). These studies suggest an involvement of SARM1-mediated innate immunity in neuronal death.

Two independent genetic screens identified SARM1 as a mediator of Wallerian axonal degeneration after acute nerve injury (Gerdts et al., 2013; Osterloh et al., 2012). SARM1 triggers Wallerian degeneration by activating the mitogen-activated protein kinase (MAPK) cascade, through nicotinamide adenine dinucleotide (NAD^+^) depletion (Essuman et al., 2017; Gerdts et al., 2015; Gerdts et al., 2016; Sasaki et al., 2016; Walker et al., 2017; Yang et al., 2015), and by stimulating Ca^2+^ influx into injured axons (Loreto et al., 2015). In addition, SARM1 mediates axon degeneration in traumatic brain injury and vincristine-induced peripheral neuropathy (Geisler et al., 2016; Henninger et al., 2016). Therefore, SARM1 represents a central element of an axonal degeneration pathway that is triggered by disparate stimuli. These findings raise the question whether SARM1 might participate to further neurodegenerative pathologies.

Prion diseases are a group of neurodegenerative disorders characterized by the deposition of PrP^Sc^, a misfolded isoform of cellular prion protein (PrP^C^), in the central nervous system (CNS) and other organs (Aguzzi and Zhu, 2012). Progressive accumulation of PrP^Sc^ in the CNS leads to lethal encephalopathies associated with neuronal loss, spongiform change and neuroinflammation (Aguzzi et al., 2013b). Prion diseases are unique among various neurodegenerative disorders since they can be acquired by infection. The contribution of innate immunity to prion pathogenesis remains to be fully understood (Aguzzi and Zhu, 2017). In the mouse scrapie model of prion disease, MyD88-mediated TLR signaling does not play a significant role in prion pathogenesis (Prinz et al., 2003). However, TLR4 signaling (probably mediated by MyD88-independent pathways) plays a protective function in prion diseases (Spinner et al., 2008).

Axonal degeneration is an early feature of most neurodegenerative disorders including prion diseases (Conforti et al., 2014). Axonal degeneration and synaptic loss are observed both in Creutzfeldt-Jakob disease and animal models of prion diseases (Ferrer, 2002; Gray et al., 2009; Guiroy et al., 1989; Jeffrey et al., 1995; Liberski and Budka, 1999; Liberski et al., 1995; Reis et al., 2015), and often precedes neuronal death. However, the mechanism of axon degeneration and its contribution to prion pathogenesis remain unknown.

In this study, we aimed to ascertain whether SARM1-mediated innate immunity and/or SARM1-dependent axon degeneration might play a role in prion pathogenesis. First, we assessed the expression of SARM1 in prion-infected mouse brains and found that SARM1 expression was decreased upon prion infection. Prion inoculation experiments revealed that disease progression was moderately accelerated by SARM1 deficiency. Analysis of the neuroinflammatory response failed to find any difference between prion-infected *SARM1*^−/−^, *SARM1*^+/−^ and their WT littermates (*SARM1*^+/+^), indicating that SARM1-mediated innate immunity does not account for the deceleration of prion progression. We then performed a genome-wide transcriptome analysis of SARM1^+/+^ and SARM1^−/−^ mouse brains with or without prion infection. Surprisingly, we found that SARM1 ablation led to an upregulated expression of the pro-apoptotic gene X-linked inhibitor of apoptosis (XIAP)-associated factor 1 (XAF1) (Liston et al., 2001), resulting in enhanced proapoptotic caspase activity and neuronal death in prion-infected *SARM1*^−/−^ mice. In vitro experiments demonstrated that the increased susceptibility of SARM1-silenced neuronal cells to apoptotic stimuli were abrogated by ablating XAF1. Hence SARM1 deficiency sensitizes mice to prion-induced neurodegeneration, probably by creating a XAF1-driven proapoptotic environment within neurons.

## Results and discussion

### Down-regulation of SARM1 expression in prion-infected mouse brains

In contrast to other MyD88 family members, SARM1 is preferentially expressed by neurons in the brain (Kim et al., 2007). To determine whether the expression of SARM1 is altered by prion infection, we performed quantitative reverse-transcription PCR (qRT-PCR) on mRNA isolated from terminally scrapie-sick wild-type (WT) C57BL/6 mice infected with the Rocky Mountain Laboratory strain of mouse-adapted scrapie prions (passage #6, therefore named RML6). For comparison, we used age-matched WT C57BL/6 mice inoculated with non-infectious brain homogenates (NBH). We found that *Sarm1* mRNA was significantly reduced in prion-infected mouse brains (Figure 1A). In contrast, other members of TLRs and adaptor molecules were either upregulated or unchanged upon prion infection (Figure 1B-D), suggesting that SARM1 is unique among TLRs associated molecules in its modulation by prion infection.

**Figure 1.**
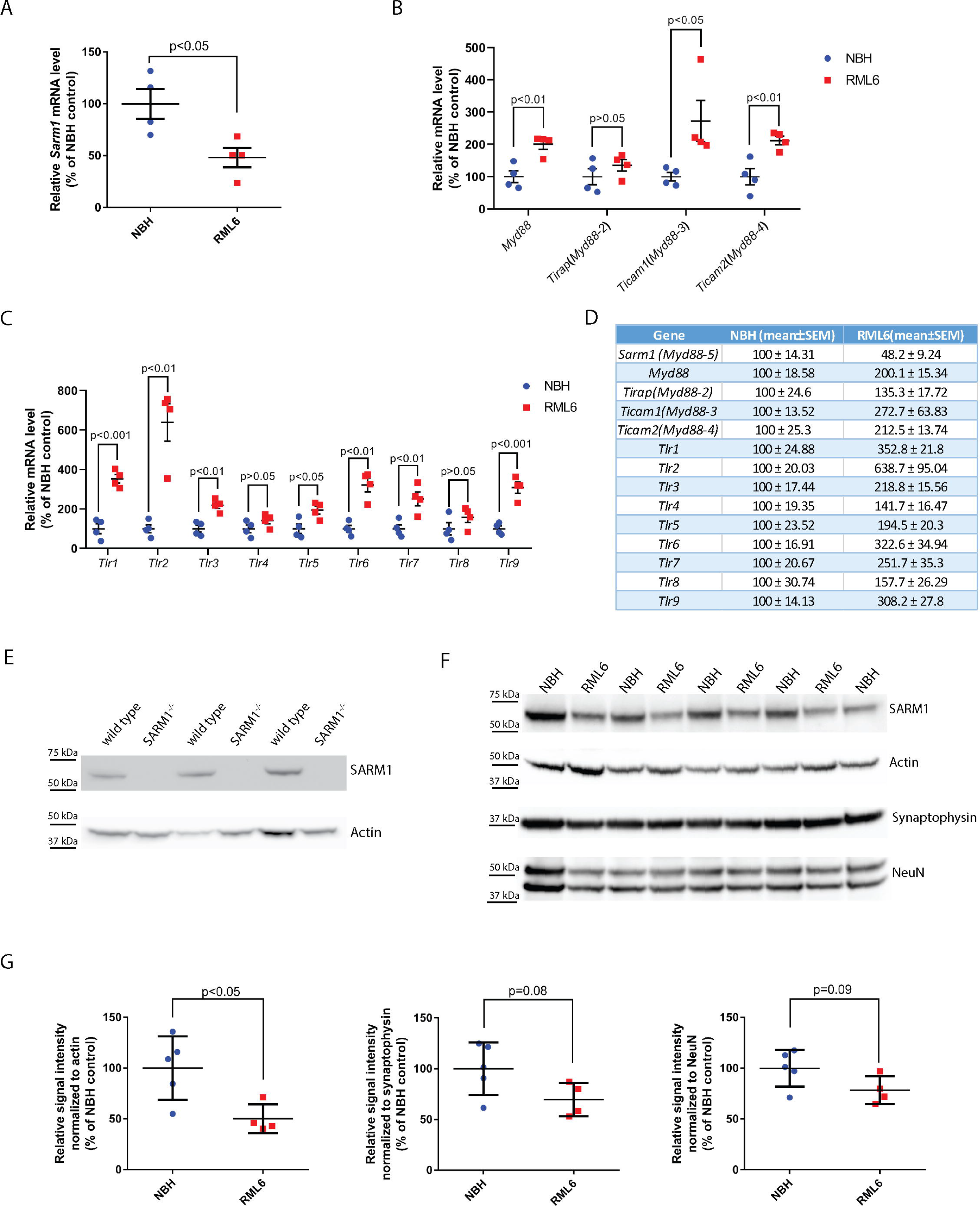
Reduced expression of SARM1 in prion-infected mouse brains. (A) qRT-PCR for *Sarm1* mRNA from terminally sick RML6-infected C57BL/6 mice and age-matched NBH-inoculated C57BL/6 mice (n=4). (B-C) qRT-PCR for MyD88 family (B) and TLRs (C) mRNA from terminally sick RML6-infected C57BL/6 mice and age-matched NBH-inoculated C57BL/6 mice (n=4). Relative expression was normalized to GAPDH expression and represented as percentage of average values in NBH-treated mice. (D) Summary of the qRT-PCR results. (B) Western blot for SARM1, Actin, Synaptophysin and NeuN on brains collected from terminally sick RML6-infected C57BL/6 mice and age-matched NBH-inoculated C57BL/6 mice. (C) Densitometric quantification of the SARM1 Western blot. n=5 for NBH-treated mice and n=4 for RML6-treated mice. Relative signal intensity was represented as percentage of average values in NBH-treated mice (n=5). qRT-PCR and Western blot results represent at least three independent experiments.

We then performed Western blot analyses to assess the SARM1 protein level in RML6 or NBH inoculated mouse brains. Again, we observed that the SARM1 protein level was significantly suppressed in RML6-infected C57BL/6 mouse brains (Figure 1E-F, full images of the cropped Western blots are shown in Supplementary Figure 1). When the SARM1 signal was normalized against neuronal markers, the relative SARM1 level was still lower in RML6 infected samples compared with NBH treated samples, indicating that the suppression of SARM1 in prion-infected mouse brains was not solely due to neuronal death at the terminal stage of disease (Figure 1F-G, full images of the cropped Western blots are shown in Supplementary Figure 1), but rather by prion-specific events (such as prion-induced stresses to the neurons).

### SARM1 deficiency resulted in accelerated prion progression

We found that SARM1 deficiency does not affect the expression of PrP^C^ in mouse brains (Figure 2A-B). To determine whether the SARM1-mediated innate immune responses and/or axonal degeneration may influence prion pathogenesis, we intracerebrally inoculated *SARM1*^−/−^, *SARM1*^+/−^ and *SARM1*^+/+^ (WT) littermates with RML6 prions. The incubation time was defined as the time lapse between prion inoculation and the time at which mice showed signs of terminal scrapie and were therefore euthanized. We found that SARM1 deficiency accelerated prion disease. *SARM1*^−/−^ mice experienced significantly shortened incubation times compared with isogenic WT littermates (median survival 166 days post inoculation (dpi) for *SARM1*^−/−^ vs. 176 dpi for WT littermates, p=0.01). *SARM1*^+/−^ mice showed a trend towards acceleration that did not reach statistical significance (median survival 171 dpi) (Figure 2C). These results indicate that SARM1 deficiency accelerates prion progression, and suggest a possible effect of SARM1 haploinsufficiency, implying that SARM1 instead plays a neuroprotective role in prion pathogenesis. The effect of SARM1 deficiency on prion progression was moderate, but it was statistically significant and its magnitude is comparable to that seen in other genetically-modified animal models or upon pharmacological treatments (Goniotaki et al., 2017; Grizenkova et al., 2014; Herrmann et al., 2015; Sakai et al., 2013; Sorce et al., 2014; Zhu et al., 2016).

**Figure 2.**
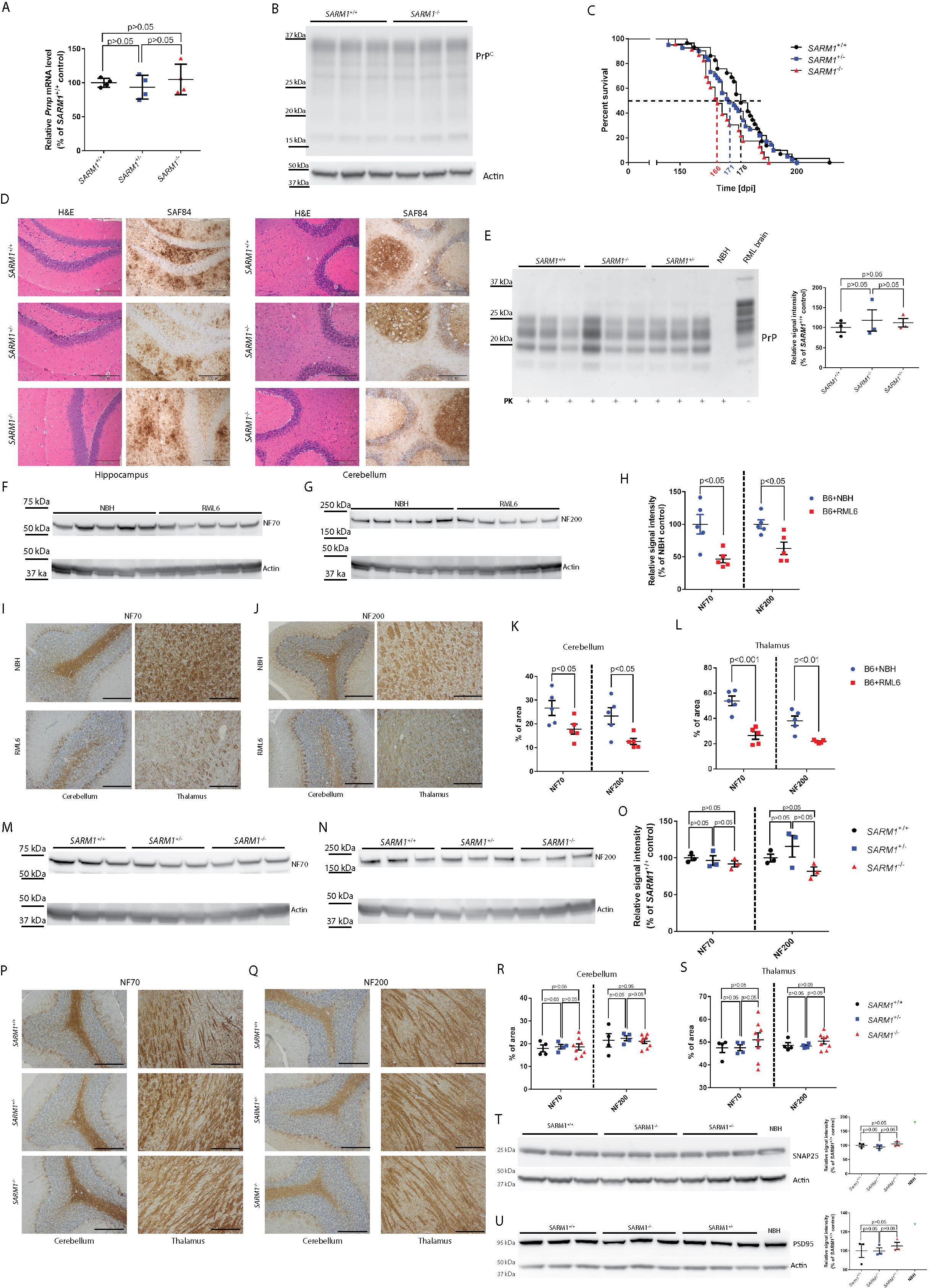
SARM1 deficiency accelerated prion progression. (A) qRT-PCR for *Prnp* mRNA from *SARM1*^−/−^, *SARM1*^+/−^ and *SARM1*^+/+^ (WT) mouse brains. n=4 for each genotype. (B) Western blot for PrP^C^ in *SARM1*^−/−^ and *SARM1*^+/+^ (WT) mouse brains. The three bands between 25 and 37 kDa identify di, mono and unglycosylated PrP^C^. (C) Survival curves of RML6 infected *SARM1*^−/−^, *SARM1*^+/−^ and *SARM1*^+/+^ (WT) mice. The median survival of *SARM1*^−/−^ mice was 166 dpi (n=23) whereas that of *SARM1*^+/+^ (WT) females was 176 dpi (n=29). *, *p*<0.05. The median survival of *SARM1*^+/−^ mice was 171 dpi (n=41). The survival curves summarize three independent i.c inoculation experiments of 22, 25 and 46 mice, respectively. (D) Hematoxylin/eosin (H&E) and immunostainings for SAF84 (which detects PrP) on RML6-infected *SARM1*^−/−^, *SARM1*^+/−^ and *SARM1*^+/+^ (WT) brains at 112 dpi. Left: hippocampus. Right: cerebellum. Scale bars: 200μm. (E) Left: PrP^Sc^ Western blot of homogenates prepared from RML6-infected *SARM1*^−/−^, *SARM1*^+/−^ and *SARM1*^+/+^ (WT) brains at 112 dpi. Samples were digested with PK as indicated and detected with POM1. Right: densitometric quantification of the PrP^Sc^ Western blot. n=3 for each genotype. (F-G) Western blot for NF70 (F) and NF200 (G) in terminally sick RML6-infected C57BL/6 mice and age-matched NBH-inoculated C57BL/6 mouse brains. (H) Densitometric quantification showing dramatic NF70 and NF200 reduction in RML6-infected brains. (I-J) NF70 (I) and NF200 (J) immunohistochemistry of brain tissues (cerebellum and thalamus) from terminally sick RML6-infected C57BL/6 mice and age-matched NBH-inoculated C57BL/6 mice. (K-L) Quantification of NF70 and NF200 staining in cerebellum (K) and thalamus (L) showed decreased NF70 and NF200 stainings in RML6-infected brains. n=5 for NBH-treated and RML6-treated mice. Relative signal intensity represents percentage of average values in NBH-treated mice. Scale bars: 200μm. (M-N) Western blot for NF70 (M) and NF200 (N) in RML6-infected *SARM1*^−/−^, *SARM1*^+/−^ and *SARM1*^+/+^ (WT) mouse brains at 112 dpi. (O) Densitometric quantification of the NF70 and NF200 Western blots showed unchanged NF70 and NF200 levels. n=3 for each genotype. Relative signal intensity was represented as percentage of average values in *SARM1*^+/+^ mice. (P-Q) NF70 (P) and NF 200 (Q) immunohistochemistry of brain tissues (cerebellum and thalamus) from RML6-infected *SARM1*^−/−^, *SARM1*^+/−^ and *SARM1*^+/+^ (WT) mice at 112 dpi. (R-S) Quantification of NF70 and NF200 staining in cerebellum (R) and thalamus (S) showed unaltered NF70 and NF200 stainings. n=4 for *SARM1*^+/−^ and *SARM1*^+/+^ (WT) mice, n=8 for *SARM1*^−/−^ mice. The immunohistochemical staining was calculated as the percentage of brown signals over the total area. Scale bars: 200μm. (T-U) Left: Western blots for SNAP25 (T) and PSD95 (U) in RML6-infected *SARM1*^−/−^, *SARM1*^+/−^ and *SARM1*^+/+^ (WT) mouse brains at 112 dpi. Right: Densitometric quantification of the SANP25 and PSD95 Western blots showed unchanged protein levels. n=3 for each genotype. Relative signal intensity was represented as percentage of average values in *SARM1*^+/+^ mice. qRT-PCR, Western blot and histology results represent at least three independent experiments. NBH: non-infectious brain homogenates.

To compare the prion pathogenesis between *SARM1*^−/−^, *SARM1*^+/−^ and *SARM1*^+/+^ mice, brains were collected synchronously from prion-infected *SARM1*^−/−^, *SARM1*^+/−^ and *SARM1*^+/+^ mice at 112dpi, a time point at which mice did not reach terminal stage. Histology of brains from *SARM1*^−/−^, *SARM1*^+/−^ and *SARM1*^+/+^ mice harvested at 112 dpi failed to reveal any obvious difference in lesion patterns and PrP^Sc^ deposition in different areas (Figure 2D). Western blots detecting proteinase K-resistant prion protein (PrP^Sc^) confirmed similar level of PrP^Sc^ accumulation in *SARM1*^−/−^, *SARM1*^+/−^ and *SARM1*^+/+^ mouse brains at 112 dpi (Figure 2E). Therefore, SARM1 deficiency neither changed prion-induced lesion pattern nor altered PrP^Sc^ accumulation.

Prion diseases are associated with axon degeneration (Ferrer, 2002; Gray et al., 2009; Guiroy et al., 1989; Jeffrey et al., 1995; Liberski and Budka, 1999; Liberski et al., 1995; Reis et al., 2015). Accordingly, we found neurofilaments NF70 and NF200 to be decreased in prion-infected mouse brains (Figure 2F-L, full images of the cropped Western blots are shown in Supplementary Figure 1). SARM1 is required for axon degeneration induced by various stimuli, and *SARM1*^−/−^ mice are protected from Wallerian degeneration, traumatic brain injury, and vincristine-induced peripheral neuropathy (Geisler et al., 2016; Gerdts et al., 2013; Henninger et al., 2016; Osterloh et al., 2012). However, in the RML mouse model of prion disease, prion pathogenesis was aggravated in *SARM1*^−/−^ mice. The acceleration of prion progression by SARM1 deficiency therefore implies that SARM1-mediated axon degeneration does not play a major role in prion pathogenesis. Consistent with this assertion, Western blot and immunohistochemistry of the neurofilaments NF70 and NF200, as well as the protein levels of pre- and post-synaptic proteins SNAP25 and PSD95 failed to show obvious difference between *SARM1*^−/−^, *SARM1*^+/−^ and *SARM1*^+/+^ mouse brains at 112 dpi (Figure 2M-U, full images of the cropped Western blots are shown in Supplementary Figure 1). This is in accordance with a previous report that slow Wallerian degeneration (Wld^S^) mice displayed unaltered prion pathogenesis (Gultner et al., 2009).

### SARM1 deficiency does not alter prion-induced neuroinflammation

SARM1 is a TLR adaptor protein that regulates the innate immune response and elicits distinct effects in *C. elegans* and mammalian cells (Carty et al., 2006; Couillault et al., 2004). Its function in mammals is still controversial since *SARM1*^−/−^ murine macrophages showed unaltered cytokine production after TLR induction (Kim et al., 2007), perhaps because of its low expression of SARM1 in myeloid cells. The innate immune response could play an important role in neurodegenerative diseases. In a *C. elegans* model of ALS, it has been reported that SARM1-mediated innate immune response is required for neurodegeneration induced by mutant TDP-43 and FUS (Veriepe et al., 2015).

Upon challenge with West-Nile virus, *SARM1*^−/−^ mice show increased viral replication in the brain stem and enhanced mortality associated with decreased microglial activation and reduced level of tumor necrosis factor α (TNFα) (Szretter et al., 2009). Therefore, we set out to assess the extent of neuroinflammation in prion-infected *SARM1*^−/−^, *SARM1*^+/−^ and *SARM1*^+/+^ mice. We performed Iba1 immunohistochemical staining to evaluate microglial activation in *SARM1*^−/−^, *SARM1*^+/−^ and *SARM1*^+/+^ mouse brains collected at 112 dpi, and observed similar level of microglial activation in all three groups (Figure 3A-B). Sholl analysis of the microglial morphology (Kongsui et al., 2014; Papageorgiou et al., 2016) showed a similar pattern of microglial activation between the three groups (Figure 3C). Western blot of Iba1 confirmed that *SARM1*^−/−^, *SARM1*^+/−^ and *SARM1*^+/+^ mouse brains experienced similar levels of microglial activation (Figure 3D, full image of the cropped Western blot are shown in Supplementary Figure 1). Additionally, we found similar number of CD68+ cells (representing activated microglia) in various regions in *SARM1*^−/−^, *SARM1*^+/−^ and *SARM1*^+/+^ mouse brains (Figure 3E). Therefore, SARM1 deficiency does not alter prion-induced microglial activation. We also carried out GFAP immunohistochemical staining on *SARM1*^−/−^, *SARM1*^+/−^ and *SARM1*^+/+^ mouse brains collected at 112dpi to assess astrogliosis. There were no overt differences between the different genotypes (Figure 3F). Western blot of GFAP confirmed that *SARM1*^−/−^, *SARM1*^+/−^ and *SARM1*^+/+^ mouse brains experienced similar levels of astrogliosis (Figure 3G, full image of the cropped Western blot are shown in Supplementary Figure 1). Furthermore, cytokine profiling showed similar levels of Tnfα, Il-1β and Il-6 mRNAs in *SARM1*^−/−^, *SARM1*^+/−^ and *SARM1*^+/+^ mouse brains at 112 dpi (Figure 3H). Therefore, SARM1 deficiency does not overtly alter prion-induced neuroinflammation, which is considered the primary innate immune response in the CNS (Aguzzi et al., 2013a).

**Figure 3.**
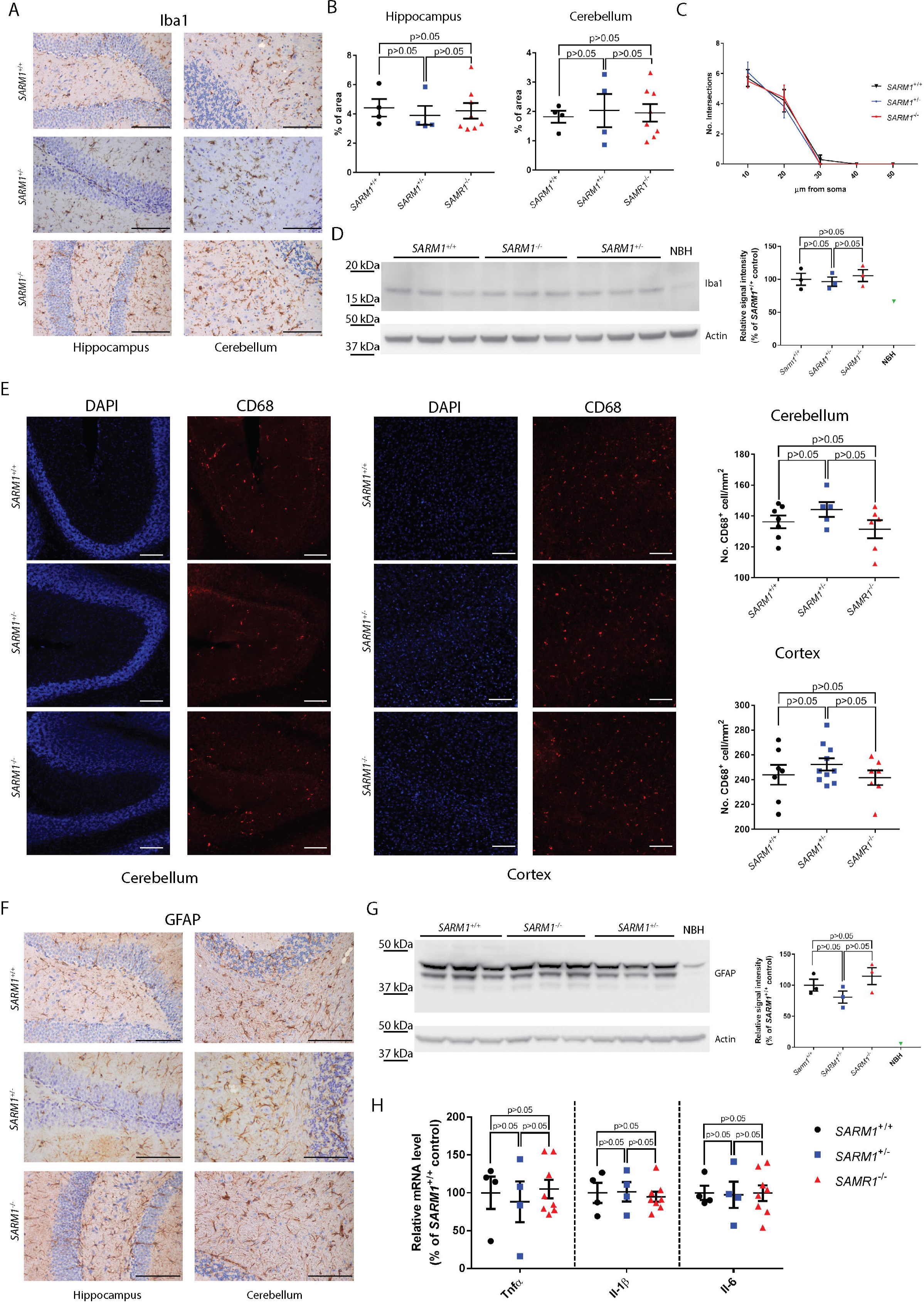
Unaltered neuroinflammation in prion-infected SARM1^−/−^ mouse brains. (A) Iba1 immunohistochemistry of brain tissues (hippocampus and cerebellum) from RML6-infected *SARM1*^−/−^, *SARM1*^+/−^ and *SARM1*^+/+^ (WT) mice at 112 dpi. Scale bars: 100μm. (B) Quantification of Iba1 staining in hippocampus (left) and cerebellum (right). n=4 for *SARM1*^+/−^ and *SARM1*^+/+^ (WT) mice, n=8 for *SARM1*^−/−^ mice. The immunohistochemical staining was calculated as the percentage of brown signals over the total area. (C) Sholl analysis of Iba1 positive microglia in RML6-infected *SARM1*^−/−^, *SARM1*^+/−^ and *SARM1*^+/+^ (WT) mouse brains at 112 dpi. (D) Left: Western blot for Iba1 in RML6-infected *SARM1*^−/−^, *SARM1*^+/−^ and *SARM1*^+/+^ (WT) mouse brains at 112 dpi. Right: densitometric quantification of the Iba1 Western blot. n=3 for each genotype. The relative signal intensity was represented as percentage of average values in *SARM1*^+/+^ mice. (E) Left: CD68 immunohistochemistry of brain tissues (left: cerebellum; right: cortex) from RML6-infected *SARM1*^−/−^, *SARM1*^+/−^ and *SARM1*^+/+^ (WT) mice at 112 dpi. Scale bars: 100μm. Right: Quantification of CD68 positive cells. n=5~10. (F) GFAP immunohistochemistry of brain tissues (left: cerebellum; right: cortex) from RML6-infected *SARM1*^−/−^, *SARM1*^+/−^ and *SARM1*^+/+^ (WT) mice at 112 dpi. Scale bars: 100μm. (G) Left: Western blot for GFAP in RML6-infected *SARM1*^−/−^, *SARM1*^+/−^ and *SARM1*^+/+^ (WT) mouse brains at 112 dpi. Right: densitometric quantification of the GFAP Western blot. n=3 for each genotype. The relative signal intensity was represented as percentage of average values in *SARM1*^+/+^ mice. (H) qRT-PCR of cytokines (Tnfα, II-1β and II-6) in RML6-infected *SARM1*^−/−^, *SARM1*^+/−^ and *SARM1*^+/+^ (WT) mouse brains at 112 dpi. n= 8 for *SARM1*^−/−^ and n=4 for *SARM1*^+/−^ and *SARM1*^+/+^ (WT) mice. Relative expression was normalized to GAPDH expression and represented as percentage of average values in *SARM1*^+/+^ mice. Histology, Western blot and qRT-PCR results represent at least three independent experiments.

The acceleration of prion pathogenesis by SARM1 ablation is consistent with findings in a West Nile virus (WNV) infection model, where *SARM1*^−/−^ mice experienced enhanced mortality. However, the pathomechanisms appear to differ between WNV and prion infections. WNV-infected *SARM1*^−/−^ mice showed decreased microglial activation and reduced levels of TNFα, whereas prion-infected *SARM1*^−/−^ mice failed to show any difference in microglial activation, astrogliosis or cytokine production.

### SARM1 deficiency leads to XAF1 upregulation and enhanced apoptosis

In an effort to discover how SARM1 deficiency accelerates prion progression, we performed whole-transcriptome analyses of *SARM1*^−/−^ and *SARM1*^+/+^ mouse brains with or without prion infection by RNAsequencing. Surprisingly, only very few genes were differentially expressed in *SARM1*^−/−^ and *SARM1*^+/+^ mouse brains. Of these, only SARM1 itself and XAF1 showed strong down- or up-regulation (fold change>2, p<0.01) in both uninfected and prion-infected brains (Figure 4A-B, Supplementary table 1). We then verified the RNAseq results by qRT-PCR to detect the mRNA level of XAF1 in *SARM1*^−/−^, *SARM1*^+/−^ and *SARM1*^+/+^ mouse brains. XAF1 mRNA was significantly upregulated by SARM1 deficiency in a gene-dose-dependent manner (Figure 4C). Interestingly, we observed that prion infection itself upregulated XAF1 expression, even in the absence of SARM1 (Figure 4C). Therefore, prion infection and SARM1 deficiency synergistically upregulated XAF1 expression in prion-infected *SARM1*^−/−^ mice.

**Figure 4.**
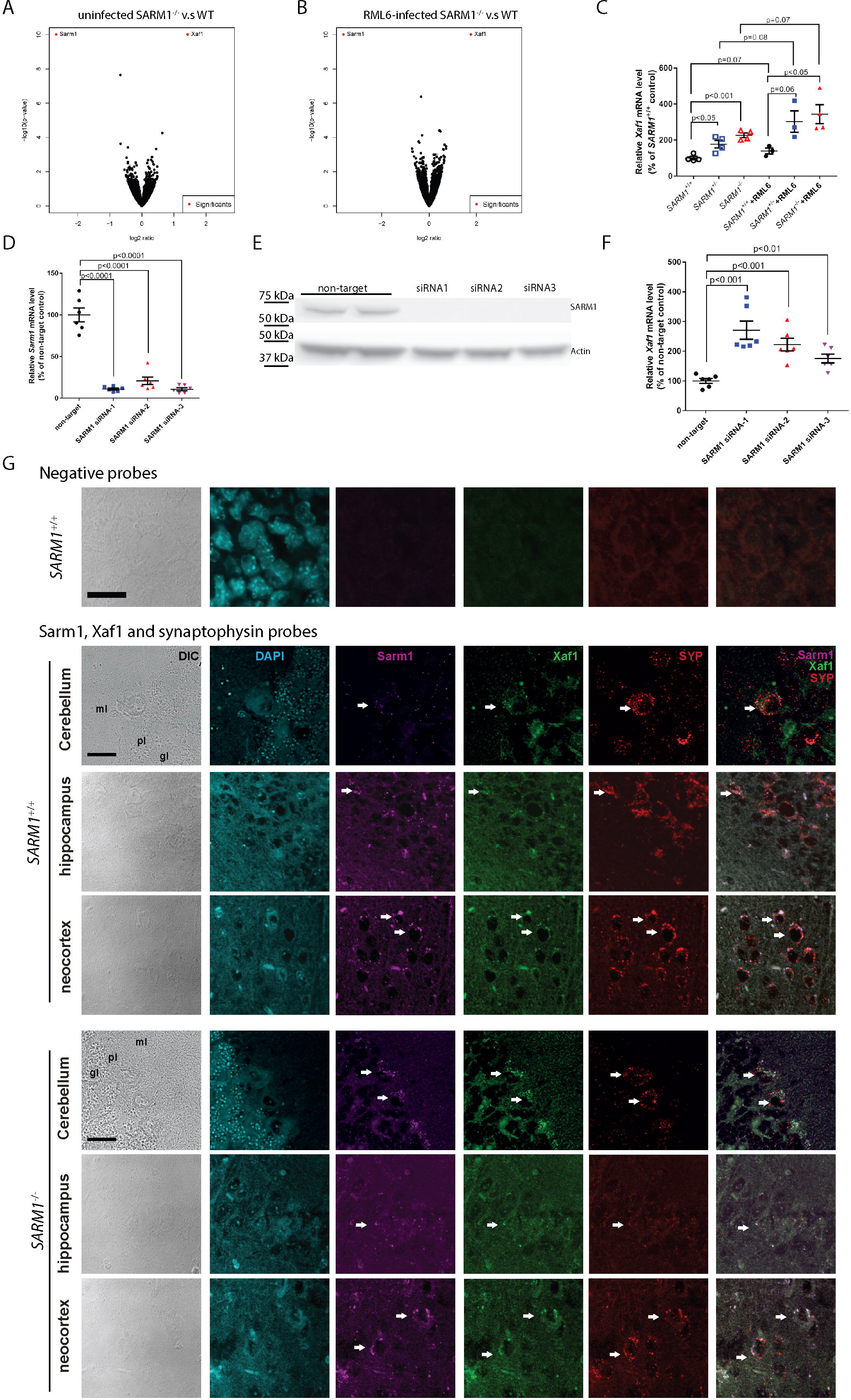
SARM1 deficiency and prion infection upregulated XAF1 expression in mouse brains. (A) Volcano plots of RNAseq data for uninfected (A) and RML6-infected (112dpi) (B) *SARM1*^−/−^ and WT mouse brains. Significants (red dots) are genes that show log2 ratio >1 or <−1 (fold change>2) with p<0.01. (C) qRT-PCR of Xaf1 in uninfected and RML6-infected (112dpi) *SARM1*^−/−^, *SARM1*^+/−^ and *SARM1*^+/+^ (WT) mouse brains. n= 3-4 for each group. Relative expression was normalized to GAPDH expression and represented as percentage of average values in uninfected *SARM1*^+/+^ mice. (D-E) qRT-PCR of Sarm1 (D) and SARM1 Western blot (E) in SARM1 siRNA treated CAD5 cells. (F) qRT-PCR of Xaf1 in SARM1 siRNA treated CAD5 cells. n=6 for each treatment. Relative expression was normalized to GAPDH expression and represented as percentage of average values in non-target control. (G) Upper panel: images of in situ hybridization using 3-Plex negative probes. Middle and lower panels: representative images of Sarm1, Xaf1 and Syp in situ hybridization in RML6-infected WT (middle) and *SARM1*^−/−^ (lower) mouse cerebella, hippocampi and neocortex at 112dpi. Differential interference contrast (DIC) and DAPI staining show the morphology and nuclei of cells. gl: granular layer; ml: molecular layer, pl: Purkinje cell layer. Lower panel: White arrows indicate Sarm1, Xaf1 and synaptophysin triple positive cells. Scale bars: 20μm. qRT-PCR and Western blot results represent at least three independent experiments. In situ hybridization represents two independent experiments.

A previous study showed upregulation of XAF1 mRNA in a different SARM1-deficient mouse line, but claimed that XAF1 protein levels remained unaltered (Hou et al., 2013). We have tested several anti-XAF1 antibodies to detect increased levels of XAF1 protein in *SARM1*^−/−^ and prion infected mouse brains. However, all tested anti-XAF1 antibodies failed to detect a specific XAF1 band. The anti-XAF1 antibody from Aviva Systems Biology (OAAB14914) could detect transiently overexpressed XAF1 in mouse neuronal CAD5 cells (XAF1 mRNA>180-fold upregulated) but not endogenous XAF1 in CAD5 cells or mouse brains (Supplementary Figure 2). Our results suggest that the failure of detecting increased level of XAF1 protein is mainly due to the lack of specific and sufficiently sensitive anti-mouse XAF1 antibodies.

The murine XAF1 and SARM1 loci are both located on chromosome 11 with a distance of 2.75 cM. The ablation of the SARM1 gene may conceivably alter XAF1 expression by affecting the chromosomal structure of neighboring loci, or by co-segregating with polymorphic DNA loci. In order to test these possibilities, we performed siRNA silencing of SARM1 in CAD5 cells. Also in this paradigm, the knockdown of SARM1 significantly upregulated XAF1 expression (Figure 4D-F, full image of the cropped Western blot are shown in Supplementary Figure 1). These results suggest that SARM1 regulates XAF1 expression in *trans*, and this regulation is cell-autonomous. The mechanisms underlying this regulation may involve miRNAs and/or epigenetic regulation (Byun et al., 2003; Lee et al., 2011; Zou et al., 2006).

To further investigate whether SARM1 deficiency leads to upregulation of XAF1 in a cell-autonomous manner in prion-infected mouse brains, we performed in situ hybridization of SARM1 and XAF1 on formalin-fixed paraffin-embedded brain sections using the RNAscope technology. Again, we found that SARM1 and XAF1 mRNA signals colocalized predominantly in synaptophysin (Syp)-positive neurons including Purkinje cells, and SARM1 deficiency increased XAF1 signals in neurons (Figure 4G). Interestingly, we also observed SARM1 signals in *SARM1*^−/−^ mouse brains in which exons 1-2 are deleted, indicating that the RNAscope SARM1 probes (20 pairs of Z RNA probe targeting exons 2-6) could still recognize the truncated SARM1 transcripts containing exons 3-6. These results confirm that the upregulation of XAF1 expression by SARM1 depletion in vivo is also cell-autonomous.

XAF1 is a pro-apoptotic molecule that antagonizes the caspase-inhibitory activity of XIAP by redistributing XIAP from the cytosol to the nucleus (Liston et al., 2001). XAF1 is expressed at low levels in many cancer cell lines, where it may contribute to immortalization and to resistance against certain apoptotic stimuli (Fong et al., 2000; Liston et al., 2001). Abolishing XAF1 expression increases the resistance of cell lines to apoptosis, whereas overexpression of XAF1 sensitizes motoneurons to axotomy and the first trimester trophoblast cells to Fas-mediated apoptosis (Perrelet et al., 2004; Straszewski-Chavez et al., 2007).

The studies discussed above indicate that XAF1 is an important mediator of cell demise. We therefore set out to assess whether SARM1 deficient mice might experience enhanced apoptosis after prion infection. We performed caspase 3 and caspase 9 activity assays by measuring their DEVDase and LEHDase activities, respectively. Indeed, RML6-infected *SARM1*^−/−^ brains showed significantly higher caspase3 and caspase 9 activity (Figure 5A-B) than SARM1^+/+^ brains collected at 112dpi. TUNEL staining showed that apoptosis was more prevalent in RML6-infected *SARM1*^−/−^ brains than in *SARM1*^+/+^ brains at 112dpi (Figure 5C). Accordingly, the neuronal marker synapsin I, which was decreased in terminally prion-sick mice (Figure 5D, full image of the cropped Western blot are shown in Supplementary Figure 1) but not altered by SARM1 deficiency in uninfected mice (Figure 5E), was significantly lower in prion-infected *SARM1*^−/−^ brains at 112dpi (5F, full image of the cropped Western blot are shown in Supplementary Figure 1). These results suggest that SARM1 deficiency sensitizes mice to prion-induced apoptosis, leading to enhanced neuronal death.

**Figure 5.**
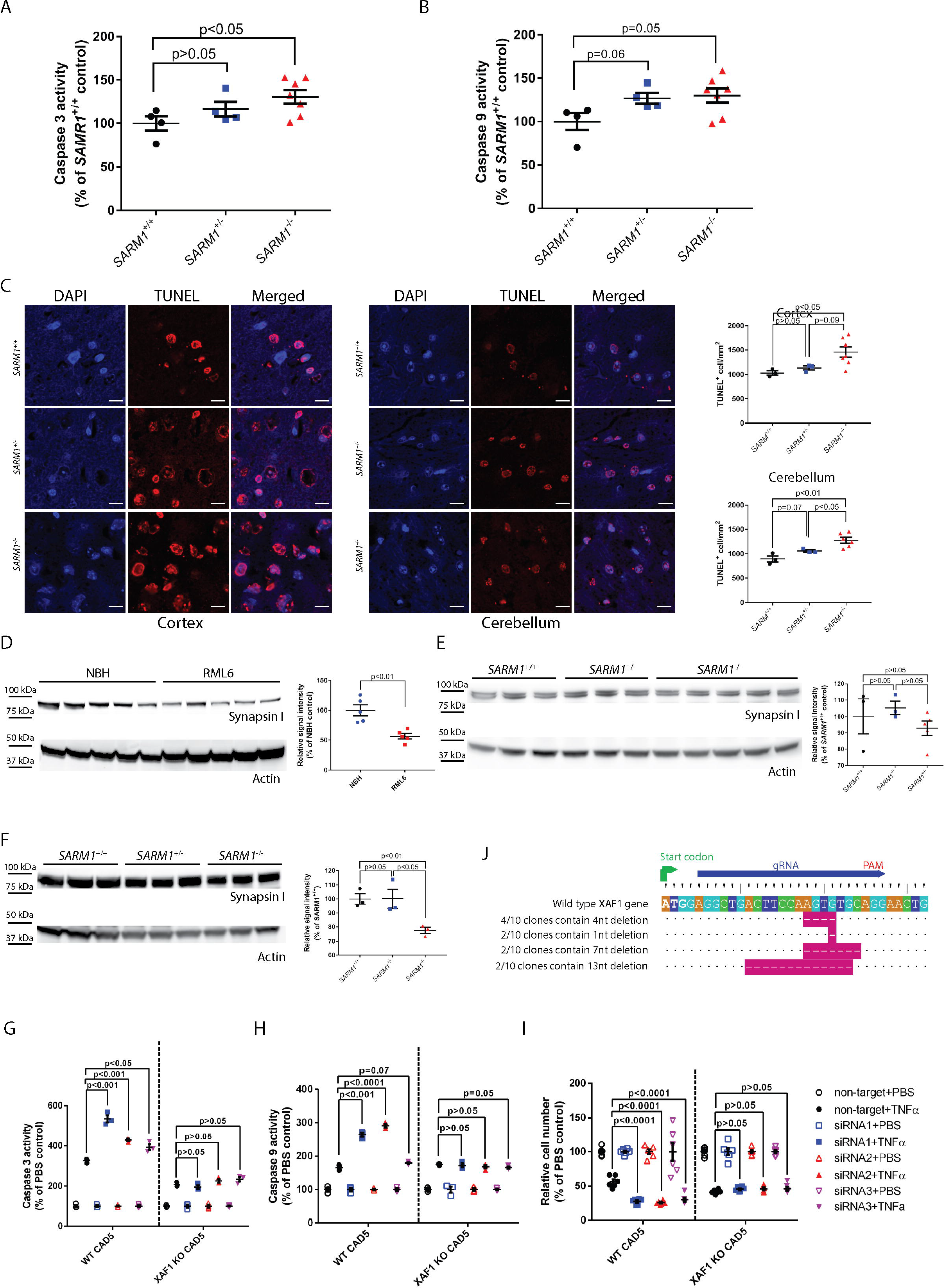
SARM1 deficiency enhanced apoptosis in prion-infected SARM1^−/−^ brains. (A-B) Caspase 3 (A) and caspase 9 (B) activity assay of RML6-infected *SARM1*^−/−^, *SARM1*^+/−^ and *SARM1*^+/+^ (WT) mouse brains at 112 dpi. n= 8 for *SARM1*^−/−^ and n=4 for *SARM1*^+/−^ and *SARM1*^+/+^ (WT) mice. Relative caspase 3 and caspase 9 activity was represented as percentage of average values in *SARM1*^+/+^ mice. (C) Left: TUNEL staining of brain tissues from RML6-infected *SARM1*^−/−^, *SARM1*^+/−^ and *SARM1*^+/+^ (WT) mice at 112 dpi. Right: Quantification of TUNEL positive cells. n=7 for *SARM1*^−/−^ and n=3 for *SARM1*^+/−^ and *SARM1*^+/+^ (WT) mice. Scale bars: 10μm. (D) Left: Western blot for synapsin I on brains collected from terminally sick RML6-infected C57BL/6 mice and age-matched NBH-inoculated C57BL/6 mice. Right: Densitometric quantification of the synapsin I Western blot showed significant reduction of synapsin I in RML6-infected brains. n=5 for NBH-treated and RML6-treated mice. Relative signal intensity was represented as percentage of average values in NBH-treated mice. (E) Left: Western blot for synapsin I in uninfected *SARM1*^−/−^, *SARM1*^+/−^ and *SARM1*^+/+^ (WT) mice. Right: Densitometric quantification of the synapsin I Western blots showed unchanged synapsin I level in uninfected *SARM1*^−/−^, *SARM1*^+/−^ and *SARM1*^+/+^ (WT) mice. n=3 for *SARM1*^+/−^ and *SARM1*^+/+^ (WT) mice, n=5 for *SARM1*^−/−^ mice. Relative signal intensity was represented as percentage of average values in *SARM1*^+/+^ mice. (F) Left: Western blot for synapsin I in RML6-infected *SARM1*^−/−^, *SARM1*^+/−^ and *SARM1*^+/+^ (WT) mouse brains at 112 dpi. Right: Densitometric quantification of the synapsin I Western blot. n=3 for each genotype. The relative signal intensity was represented as percentage of average values in *SARM1*^+/+^ mice. (G) Caspase 3 activity assay of wild type (WT) CAD5 and XAF1 KO CAD5 cell lysates after SARM1 siRNAs and TNFα treatments. (H) Caspase 9 activity assay of WT CAD5 and XAF1 KO CAD5 cell lysates after SARM1 siRNAs and TNFα treatments. n=3. Relative caspase 3 and caspase 9 activity was represented as percentage of average values in PBS groups. (I) RealTime-Glo assay of live WT CAD5 and XAF1 KO CAD5 cells after SARM1 siRNAs and TNFα treatments. n=6. Relative cell numbers were represented as percentage of average values in PBS groups. Relative caspase 3 and caspase 9 activity measurement represents two independent experiments. (J) Sequencing results of Xaf1 gene in XAF1 KO CAD5 cells showed that the coding sequence of all sequenced clones was frameshifted. Western blot results represent at least three independent experiments.

It has been reported that XAF1 may cooperate with TNFα to induce apoptosis and increased expression of XAF1 sensitized cells to apoptosis (Straszewski-Chavez et al., 2007; Xia et al., 2006). To establish whether the enhanced susceptibility to apoptosis by SARM1 deficiency is due to the upregulation of XAF1, we first silenced SARM1 in CAD5 cells and then treated them with TNFα. We found that while TNFα led to apoptosis and reduced cell numbers in both SARM1-silenced and non-target siRNA treated cells, apoptosis was enhanced and cell numbers were more conspicuously reduced in SARM1-silenced cells (Figure 5G-I). However, when the same treatment was performed on XAF1 KO CAD5 cells generated by CRISPR/Cas9 system (Figure 5J), the enhancement by SARM1 silencing was abrogated (Figure 5G-I), indicating that SARM1 deficiency sensitized cells to apoptotic stimuli mainly through the upregulated XAF1. Therefore, it is conceivable that the deterioration of prion pathogenesis in *SARM1*^−/−^ mice may be due to the upregulated XAF1 and consequently enhanced apoptosis.

In summary, we found that SARM1 deficiency unexpectedly accelerated prion progression. SARM1 deficiency selectively upregulated expression of the pro-apoptotic protein XAF1, which in turn enhanced apoptosis in prion-infected *SARM1*^−/−^ mice. Since the effects appear to be gene-dosage dependent, XAF1 repression may potentially ameliorate prion-related neuronal pathologies.

## Methods

### Ethical statement

All animal experiments were carried out in strict accordance with the Rules and Regulations for the Protection of Animal Rights (Tierschutzgesetz and Tierschutzverordnung) of the Swiss Bundesamt für Lebensmittelsicherheit und Veterinärwesen and were preemptively approved by the Animal Welfare Committee of the Canton of Zürich (permit 41/2012).

### Animals

*SARM1*^−/−^ mice (Szretter et al., 2009) carrying a targeted deletion of exons 1-2 were first backcrossed to C57BL/6J to obtain *SARM1*^+/−^ offspring. *SARM*^+/−^ mice were then intercrossed to obtain *SARM*^−/−^, *SARM1*^+/−^ and *SARM*^+/+^ (WT) littermates, which were used for the experiments described here. Mice were maintained in high hygienic grade facility and housed in groups of 3-5, under a 12 h light/12 h dark cycle (from 7 am to 7 pm) at 21±1°C, with sterilized food (Kliba No. 3436, Provimi Kliba, Kaiseraugst, Switzerland) and water *ad libitum*.

### Intracerebral prion inoculation

Mice at 6-8 weeks old were intracerebrally (i.c) inoculated with 30 μl of brain homogenate diluted in PBS with 5% BSA and containing 3 × 10^5^ LD50 units of the RML6. Scrapie was diagnosed according to clinical criteria (ataxia, kyphosis, priapism, and hind leg paresis). Mice were sacrificed on the day of onset of terminal clinical signs of scrapie.

### Quantitative real-time PCR (qRT-PCR)

Total RNA from brain was extracted using TRIzol (Invitrogen Life Technologies) according to the manufacturer’s instruction. The quality of RNA was analyzed by Bioanalyzer 2100 (Agilent Technologies), RNAs with RNA integrity number >7 were used for cDNA synthesis. cDNA were synthesized from ~1 μg total RNA using QuantiTect Reverse Transcription kit (QIAGEN) according to the manufacturer’s instruction. Quantitative real-time PCR (qRT-PCR) was performed using the SYBR Green PCR Master Mix (Roche) on a ViiA7 Real-Time PCR system (Applied Biosystems). The following primer pairs were used: GAPDH sense 5′-CCA CCC CAG CAA GGA GAC T-3′; antisense, 5′-GAA ATT GTG AGG GAG ATG CT-3′. Sarm1(Myd88-5) sense, 5′-CGC TGC CCTGTA CTG GAG G-3′; antisense, 5′-CTT CAG GAG GCT GGC CAG CT-3′. Myd88 sense, 5’-TCA TGT TCT CCA TAC CCT TGG T-3’; antisense, 5’-AAA CTG CGA GTG GGG TCA G-3’. Tirap (Myd88-2) sense, 5’-CCT CCT CCA CTC CGT CCA A-3’; antisense, 5’-CTT TCC TGG GAG ATC GGC AT-3’. Ticam1 (Myd88-3) sense, 5’-CCA GCT CAA GAC CCC TAC AG-3’; antisense, 5’-CAA GGC ACC TAG AAT GCC AAA-3’;. Ticam2 (Myd88-4) sense, 5’-CGA TCA AGA CGG CCA TGA GTC-3’; antisense, 5’-CTC GTC GGT GTC ATC TTC TGC-3’. Tlr1 sense, 5’-TGA GGG TCC TGA TAA TGT CCT AC-3’; antisense, 5’-AGA GGT CCA AAT GCT TGA GGC-3’. Tlr2 sense, 5’-GCA AAC GCT GTT CTG CTC AG-3’; antisense, 5’-AGG CGT CTC CCT CTA TTG TAT T-3’. Tlr3 sense, 5’-GTG AGA TAC AAC GTA GCT GAC TG-3’; antisense, 5’-TCC TGC ATC CAA GAT AGC AAG T-3’. Tlr4 sense, 5’-AGG CAC ATG CTC TAG CAC TAA-3’; antisense, 5’-AGG CTC CCC AGT TTA ACT CTG-3’. Tlr5 sense, 5’-GCA GGA TCA TGG CAT GTC AAC-3’; antisense, 5’-ATC TGG GTG AGG TTA CAG CCT-3’. Tlr6 sense, 5’-TGA GCC AAG ACA GAA AAC CCA-3’; antisense, 5’-GGG ACA TGA GTA AGG TTC CTG TT-3’. Tlr7 sense, 5’-ATG TGG ACA CGG AAG AGA CAA-3’; antisense, 5’-GGT AAG GGT AAG ATT GGT GGT G-3’. Tlr8 sense, 5’-GAA AAC ATG CCC CCT CAG TCA-3’; antisense, 5’-CGT CAC AAG GAT AGC TTC TGG AA-3’. Tlr9 sense, 5’-ATG GTT CTC CGT CGA AGG ACT-3’; antisense, 5’-GAG GCT TCA GCT CAC AGG G-3’. Tnfα sense, 5′-CAT CTT CTC AAA ATT CGA GTG ACA A-3′; antisense, 5′-TGG GAG TAG ACA AGG TAC AAC CC-3′. II-1β sense, 5′-CAA CCA ACA AGT GAT ATT CTC CAT G-3′; antisense, 5′-GAT CCA CAC TCT CCA GCT GCA-3′. Il-6 sense, 5′-TCC AAT GCT CTC CTA ACA GAT AAG-3′; antisense, 5′-CAA GAT GAA TTG GAT GGT CTT G-3′. Xaf1 sense, 5′-GAA GCT TGA CCA TGG AGG CT-3′; antisense, 5′-GGT GCA CAA CTT CCA TGT GCT-3′. Expression levels were normalized using GAPDH.

### Western blot analysis

To detect SARM1 and PrP^C^ in the mouse brains, one hemisphere from each brain was homogenized with buffer PBS containing 0.5% Nonidet P-40 and 0.5% CHAPSO. Total protein concentration was determined using the bicinchoninic acid assay (Pierce). ~20 μg protein were loaded and separated on a 12% Bis-Tris polyacrylamide gel (NuPAGE, Invitrogen) and then blotted onto a nitrocellulose membrane. Membranes were blocked with 5% wt/vol Topblock (LuBioScience) in PBS supplemented with 0.05% Tween 20 (vol/vol) and incubated with primary antibodies anti-SARM1 (D2M5I) rabbit mAb (1:1000; Cell Signaling Technology, 13022) or POM1 (200 ng ml^−1^) in 1% Topblock overnight. After washing, the membranes were then incubated with secondary antibody horseradish peroxidase (HRP)-conjugated goat anti-rabbit IgG (1:10,000, Jackson ImmunoResearch, 111-035-045) or goat anti–mouse IgG (1:10,000, Jackson ImmunoResearch, 115-035-003). Blots were developed using Luminata Crescendo Western HRP substrate (Millipore) and visualized using the Stella system (model 3200, Raytest.). To avoid variation in loading, the same blots were stripped and incubated with an anti-actin antibody (1:10,000, Millipore, MAB1501R). The signals were normalized to actin as a loading control. To detect PrP^Sc^ in prion infected *SARM1*^−/−^, *SARM1*^+/−^ and *SARM1*^+/+^ (WT) mouse brains, prion infected forebrains were homogenized in sterile 0.32 M sucrose in PBS. Total protein concentration was determined using the bicinchoninic acid assay (Pierce). Samples were adjusted to 20 μg protein in 20 μl and digested with 25 μg ml^−1^ proteinase K in digestion buffer (PBS containing 0.5% wt/vol sodium deoxycholate and 0.5% vol/vol Nonidet P-40) for 30 min at 37°C. PK digestion was stopped by adding loading buffer (Invitrogen) and boiling samples at 95°C for 5 min. Proteins were then separated on a 12% Bis-Tris polyacrylamide gel (NuPAGE, Invitrogen) and blotted onto a nitrocellulose membrane. POM1 and horseradish peroxidase (HRP)-conjugated goat anti–mouse IgG were used as primary and secondary antibodies, respectively. Blots were developed using Luminata Crescendo Western HRP substrate (Millipore) and visualized using the FUJIFILM LAS-3000 system. To detect NeuN, synaptophysin, NF70, NF200, SNAP25, PSD95, Iba-1, GFAP and synapsin I in prion-infected wild type C57BL/6 mouse brains or *SARM1*^−/−^, *SARM1*^+/−^ and *SARM1*^+/+^ (WT) mouse brains, 20 μg of total brain protein were loaded and anti-NeuN clone EPR12763 (Abcam, ab177487), anti-synaptophysin Clone 2/synaptophysin (BD Biosciences, 611880), anti-neurofilament-L clone DA2 (1:1000, Cell Signaling Technology, 2835), anti-neurofilament 200 clone NE14 (1:1000, Sigma, N5389), anti-SNAP25 antibody (1:1000; Abcam, ab5666), anti-PSD95 antibody (1:1000; Abcam, 18258), anti-Iba-1 antibody (1:1000; Wako Chemicals GmbH, Germany), anti-GFAP antibody (D1F4Q) XP^®^ Rabbit mAb (1:3000; Cell Signaling Technology, 12389) and anti-synapsin I (1:2000, Millipore AG, AB1543) were used. To detect XAF1 in mouse brains and CAD5 cells, primary antibodies rabbit anti–mouse XAF1 antibody (1:1000; Abcam, ab17204 and ab81353; Aviva Systems Biology, OAAB14914; Boster, A03432; Invitrogen, PA1-41099; GeneTex, GTX51339; Novus Biologicals, NB100-56355) were used. Actin was used as loading control.

### Immunohistochemistry

For immunohistochemistry of prion-infected brains, formalin-fixed tissues were treated with concentrated formic acid for 60 min to inactivate prion infectivity and embedded in paraffin. Paraffin sections (2 μm) of brains were stained with hematoxylin/eosin (H&E). After deparaffinization through graded alcohols, Iba-1 antibody (1:1000; Wako Chemicals GmbH, Germany) was used for highlighting microglial cells, GFAP antibody (1:300; DAKO, Carpinteria, CA) was used for astrocytes, anti-neurofilament 70 Clone 2F11 (1:100, DAKO, M076229-2) and anti-neurofilament 200 clone NE14 (1:2000, Sigma, N5389) antibodies were used for assessing axonal degeneration. Stainings were visualized using an IVIEW DAB Detection Kit (Ventana), with a hematoxylin counterstain applied subsequently. For the histological detection of partially proteinase K-resistant prion protein deposition, deparaffinized sections were pretreated with formaldehyde for 30min and with 98% formic acid for 6 min, and then washed in distilled water for 30 min. Sections were incubated in Ventana buffer and stains were performed on a NEXEX immunohistochemistry robot (Ventana instruments, Switzerland) using an IVIEW DAB Detection Kit (Ventana). After incubation with protease 1 (Ventana) for 16 min, sections were incubated with anti-PrP SAF-84 (SPI bio, A03208, 1:200) for 32 min. Sections were counterstained with hematoxylin. Sections were imaged using a Zeiss Axiophot light microscope. Quantification of Iba-1, GFAP, NF70 and NF200 staining was performed on acquired images, where regions of interest were drawn, and the percentage of brown Iba-1, GFAP, NF70 and NF200 staining over the total area was quantified using Image J software (National Institutes of Health). For CD68 staining, 4% PFA fixed brain tissues were cut into 10μm cryosections and stained with anti-CD68 antibody clone FA-11 (Bio-Rad, MCA1957) and visualized by Alexa 546 conjugated goat-anti-rat antibody (Invitrogen, A11081). The nuclei were counterstained with DAPI. Images were captured using confocal laser scanning microscope FluoView Fv10i (Olympus). The operator was blind to the genotype and treatment of the analyzed tissues.

### RNAsequencing

Total RNA from brain was extracted using TRIzol (Invitrogen Life Technologies) according to the manufacturer’s instruction, followed by RNA cleanup using RNeasy kit (Qiagen). The quality of RNA was analyzed by Bioanalyzer 2100 (Agilent Technologies), RNAs with RNA integrity number >8 were further processed. The TruSeq Stranded mRNA Sample Prep Kit (Illumina, Inc, California, USA) was used in the succeeding steps. Briefly, total RNA samples (500 ng) were poly-A selected and then reverse-transcribed into double-stranded cDNA with Actinomycin added during first-strand synthesis. The cDNA samples was fragmented, end-repaired and adenylated before ligation of TruSeq adapters. The adapters contain the index for multiplexing. Fragments containing TruSeq adapters on both ends were selectively enriched with PCR. The quality and quantity of the enriched libraries were validated using Qubit^®^ (1.0) Fluorometer (Life Technologies, California, USA) and the Bioanalyzer 2100 (Agilent Technologies). The product is a smear with an average fragment size of approximately 360 bp. The libraries were normalized to 10nM in Tris-Cl 10 mM, pH8.5 with 0.1% Tween 20. The TruSeq SR Cluster Kit v4-cBot-HS (Illumina, Inc, California, USA) was used for cluster generation using 8 pM of pooled normalized libraries on the cBOT. Sequencing were performed on the Illumina HiSeq 4000 single end 125 bp using the TruSeq SBS Kit v4-HS (Illumina, Inc, California, USA). Reads were quality-checked with FastQC. For RNAseq data analysis, we used RSEM to perform RNAseq read alignment and expression estimation (Li and Dewey, 2011). As reference we used the GRCm38 genome assembly and the corresponding gene annotations provided by Ensembl. We computed differential expression with the Bioconductor package DESeq2 (Love et al., 2014). A gene was considered as differentially expressed by applying a threshold of 0.01 for the p-value and 1 for the log-ratio. Additionally, we filtered away genes that had very low counts. Specifically we did not consider a gene as expressed if it did not exceed in at least one condition an average read count of 10 in the samples. All clusterings and visualizations were performed with R/Bioconductor (R version 3.2).

### Establishment of XAF1 KO CAD5 cells by CRISPR/Cas9

A XAF1 knockout CAD5 cell line was established by CRISPR/Cas9 genome editing approach. Briefly, a guide RNA (gRNA) targeting exon 1 of the XAF1 gene (5′-AGGCTGACTTCCAAGTGTGC-3′) was cloned into the pKLV-U6gRNA(BbsI)-PGKpuro2ABFP vector (Addgene 50946) to generate pKLV-XAF1 sgRNA plasmid. CAD5 cells were co-transfected with pKLV-XAF1 sgRNA and pCMV-hCas9 (Addgene 41815) in a molar ratio 1:1 by Lipofectamine 2000 (Invitrogen, 11668-019). Transfected cells were selected and enriched by puromycin (2μg/ml) and single cell clones were obtained by limited dilution. Cell clones were grown and expanded. To confirm gene editing of the XAF1 gene in cell clones, a region embracing exon 1 of the XAF1 gene was PCR-amplified using a forward primer (5′-GTC ACA GTC ACG GTA GCA CA-3′) and reverse primer (5′- GGG ACC AGT CCC TCC ATA CT-3′), the PCR product was cloned into pCR-Blunt II-TOPO vector (Invitrogen, 45-0245) and sequenced to verify the frameshifted indels.

### siRNA and TNFα treatment on CAD5 and XAF1 KO CAD5 cells

24 hours before transfection, 2 × 10^5^ CAD5 cells (murine neuronal cell line) per well were seeded in 6-well plates. CAD5 cell were then transfected with three different SARM1 siRNA (Thermo Fisher Scientific, siRNA ID: s108645, s108646 and s108647) or non-target negative control siRNA (Thermo Fisher Scientific, 4390843) at 50nM concentration using Lipofectamine^®^ RNAiMAX. 48 hours after transfection, cells were harvested for total RNA extraction and protein analysis.

For TNFα treatment, 2 × 10^4^ CAD5 or XAF1 KO CAD5 cells per well were first seeded in 96-well plates and transfected with SARM1 siRNAs or non-target negative control siRNA. 24 hours after the transfection, cells were treated with TNFα at a concentration 50ng/ml or PBS control for 72 hours. Cell viability was assessed by RealTime-Glo (Promage G9713). The relative cell viability was calculated as percentages to the PBS control groups.

### RNAscope in situ hybridization

Transcripts of SARM1, XAF1 and Syp in formalin-fixed paraffin-embedded brain sections were detected by in situ hybridization using RNAscope Multiplex Fluorescent Reagent Kit v2 (Advanced Cell Diagnostics, 323100) following manufacturer’s manual. Briefly, sections were deparaffinized and pretreated with hydrogen peroxide and protease plus, followed by hybridization with target probes (SARM1, 504191-C2; XAF1, 504181; Syp, 426521-C3) which are designed and manufactured by Advanced Cell Diagnostics. 3-Plex negative control probes targeting DapB gene from the Bacillus subtilis strain SMY (320871) were used as negative control. After amplification steps, fluorescence signals were developed by TSA Plus fluorophores kits (PerkinElmer, fluorescein, NEL741E001KT; Cyanine 3, NEL744E001KT; Cyanine 5, NEL745001KT). Images were captured using Leica TCS SP5 confocal microscope (Leica Microsystem).

### Caspase 3 and Caspase 9 activity assay

Caspase 3 activity was measured by Caspase 3 assay kit (Abcam, ab39401) and Caspase 9 activity was measured by Caspase 9 assay kit (Abcam, ab65608) according to the manufacturer’s instruction. Briefly, brain homogenates or cell lysates containing the same amount of total protein were incubated with either DEVE-p-NA or LEHD-p-NA substrates. After cleavage, the p-NA light emission was quantified using a microtiter plate reader at 405 nm. The activity of Caspase 3 and Caspase 9 was calculated as percentages to the SARM1^+/+^ (wt) control or cell lysates treated with PBS.

### TUNEL assay

Apoptosis in paraffin sections of prion-infected mouse brains were assesses by In situ BrdU-Red DNA fragmentation assay kit according to the manufacturer’s instructions (Abcam, ab66110). Briefly, sections were deparaffinized and treated with proteinase K, followed by labeling of DNA nicks with Br-dUTP (bromolated deoxyuridine triphosphate nucleotide) mediated by TdT (terminal deoxynucleotidyl transferase). The incorporated BrdU was detected by fluorescently labeled anti-BrdU monoclonal antibody. The nuclei were counterstained with DAPI. Images were captured using confocal laser scanning microscope FluoView Fv10i (Olympus) and analyzed using Image J software (National Institutes of Health).

### Statistical analysis

Results are presented as the mean of replicas ± SEM. Unpaired Student’s t-test was used for comparing two samples. For *in vivo* experiments, all groups were compared by Log-rank (Mantel-Cox) test. *p*-values <0.05 were considered statistically significant.

## Supporting information

## Acknowledgements

We thank the team of the Institute of Neuropathology, University Hospital Zurich, and in particular I. Abakumova, R. Moos, J. Guo, M. Delic, K. Arroyo, and M. König for technical assistance. We thank Elisabeth J Rushing for reading and editing the manuscript. We also thank Marco Colonna from Washington University School of Medicine for providing us with *SARM1*^−/−^ mice.

A. Aguzzi is the recipient of an Advanced Grant of the European Research Council (ERC, No. 250356) and is supported by grants from the European Union (PRIORITY, NEURINOX), the GELU foundation, the Swiss National Foundation (SNF, including a Sinergia grant), the Swiss Initiative in Systems Biology, SystemsX.ch (PrionX, SynucleiX), and the Klinische Forschungsschwerpunkte (KFSPs) "small RNAs" and "Human Hemato-Lymphatic Diseases". K. Frontzek is a recipient of a grant from the Theodor und Ida Herzog-Egli Stiftung. The funders had no role in study design, data collection and analysis, decision to publish, or preparation of the manuscript.

The authors declare no competing financial interests.

Author contributions: C.Zhu and A. Aguzzi conceived and designed the overall study. C. Zhu and B. Li performed most of the experiments and interpreted the data. K. Frontzek contributed to the generation of the XAF1 KO CAD5 cell line and captured in situ hybridization images. Y. Liu performed the RNAscope in situ hybridization. C.Zhu, B.Li and A.Aguzzi wrote the manuscript.

---

***Supplementary figure 1***

Full images of the cropped Western blots in figure 1E, 1F, 2F, 2G, 2M, 2N, 2T, 2U, 3D, 3G, 4E, 5D, 5E and 5F.

***Supplementary figure 2***

(A-G) Western blots for XAF1 in CAD5 cell lysates using various anti-XAF1 antibodies: (A) Abcam (ab17204); (B) Abcam (ab81353); (C) Aviva System Biology (OAAB14914); (D) Boster (A03432); (E) GeneTex (GTX51339); (F) Invitrogene (PA1-41099); (G) Novus Biologicals (NB100-56355). Lane 1: 20ug of untransfected CAD5 cell lysates; labe 2: 20ug of CAD5 transfected with pcDNA-XAF1 cell lysates; lane 3: 20ug of CAD5 transfected with pcDNA empty vector cell lysates. Arrow: specific XAF1 band; Asterisk: unspecific bands. (H) qRT-PCR for Xaf1 in CAD5 cells transfected with pcDNA empty vector or pcDNA-XAF1 plasmids. Relative expression normalized to GAPDH expression and represented as fold change compared to pcDNA transfected CAD5 cells. (I) Western blots for XAF1 in CAD5 cell lysates using Aviva System Biology (OAAB14914) anti-XAF1 antibodies. Lane 4: 60ug of wild type CAD5 cell lysates; lane 5: 60ug of XAF1 KO CAD5 cell lysates; lane 6: 20ug of CAD5 transfected with pcDNA empty vector cell lysates; lane 7: 20ug of CAD5 transfected with pcDNA-XAF1 plasmids cell lysates. Only the XAF1 antibody from Aviva Systems Biology (OAAB14914) could detect transiently overexpressed XAF1 in CAD5 cells but not endogenous XAF1 in CAD5 cells or mouse brains. Arrow: specific XAF1 band; Asterisk: unspecific bands. (J-P) Western blots for XAF1 in brain homogenates from *SARM1*^−/−^, and *SARM1*^+/+^ (WT) mice using various anti-XAF1 antibodies: (J) Abcam (ab17204); (K) Abcam (ab81353); (L) Aviva System Biology (OAAB14914); (M) Boster (A03432); (N) GeneTex (GTX51339); (O) Invitrogene (PA1-41099); (P) Novus Biologicals (NB100-56355). 20ug of *SARM1*^+/+^ (WT) or *SARM1*^−/−^ brain homogenates were loaded. Asterisk: unspecific bands with wrong sizes. Hollow arrow: unspecific band with expected size.

***Supplementary table 1***

Differentially expressed genes (DEGs) between uninfected *SARM1*^−/−^ and *SARM1*^+/+^ (WT) mouse brains (Table 1A) and RML6-infected *SARM1*^−/−^ and *SARM1*^+/+^ (WT) mouse brains (Table 1B).

## Abbreviations

CNS: central nervous system
dpi: days post inoculation
MyD88: myeloid differentiation primary response gene 88
NBH: non-infectious brain homogenates
PK: proteinase K
qRT-PCR: quantitative real-time PCR
RML6: Rocky Mountain Laboratories scrapie strain passage 6
SARM1: sterile α HEAT/armadillo motifs containing protein
XAF1: X-linked inhibitor of apoptosis-associated factor 1

