## Supplementary figures and images for "SARM1 deficiency, which prevents Wallerian degeneration, upregulates XAF1 and accelerates prion disease"

### Supplementary file 1

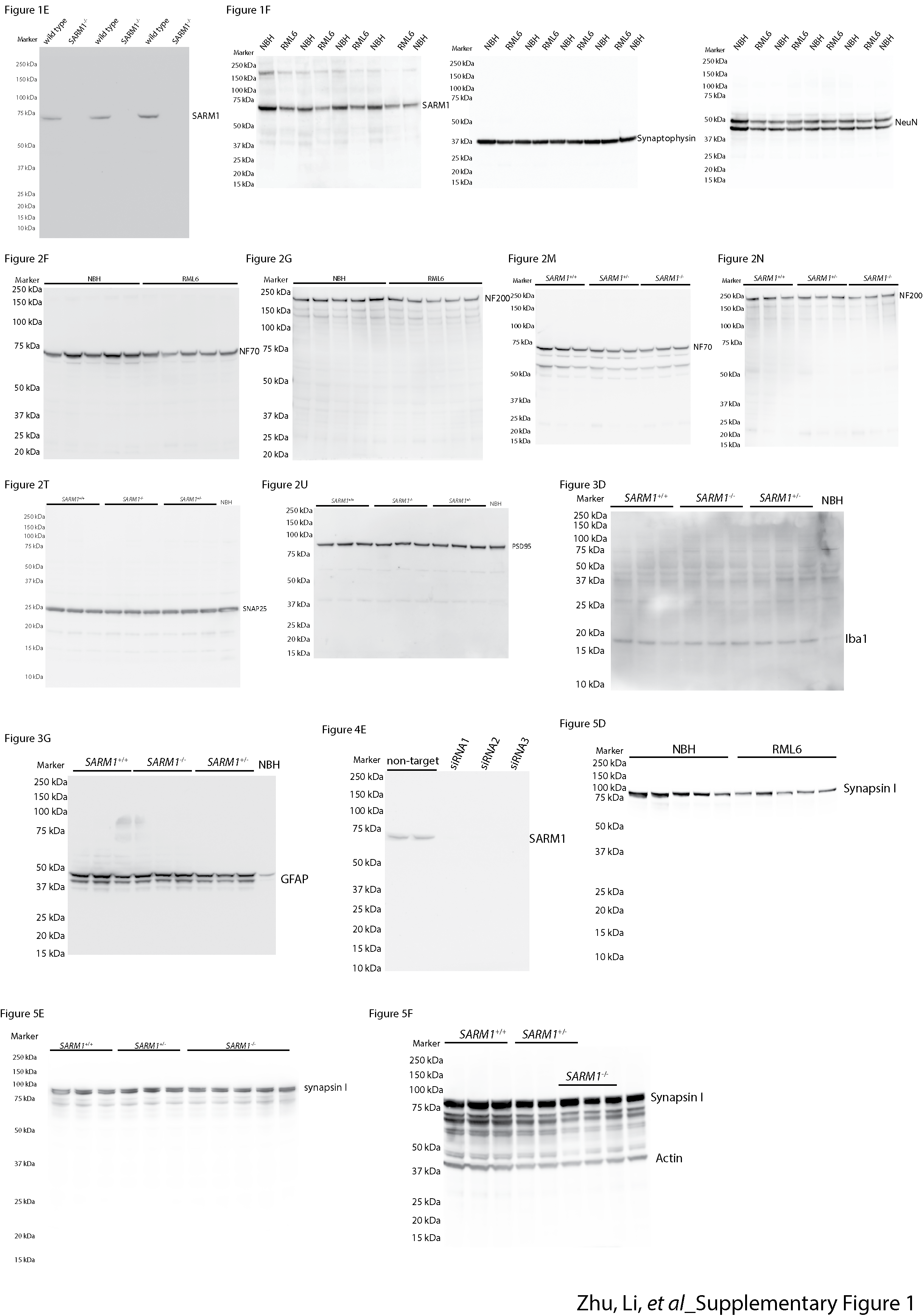

### Supplementary file 2

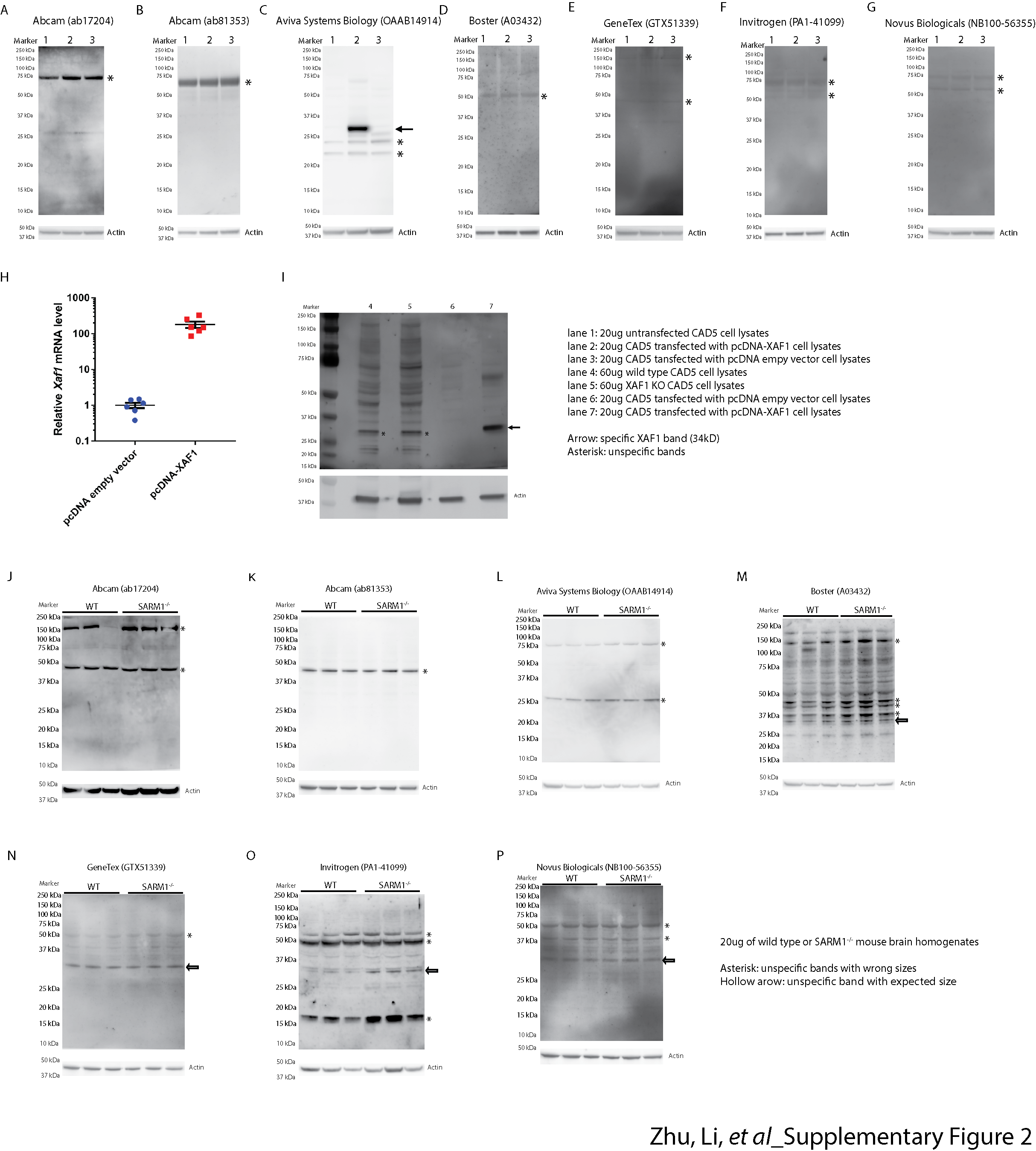
